# Sex-specific associations of estradiol and instrumental harm with moral decision-making

**DOI:** 10.64898/2026.07.08.737159

**Authors:** Aistė Ambrasė, Melina Grahlow, Erik Ilkevič, Ximena Lenz, Niklas Klink, Ramunė Grikšienė, Birgit Derntl

## Abstract

Previous research suggests that women and men may differ in moral decision-making involving instrumental harm. However, prior findings have yielded inconsistent explanations for these differences, suggesting effects of gender expression, empathy, action tendencies, and biological causes, such as differences in gonadal hormone levels. Yet, no data exists on the effects of endogenous estradiol and progesterone on moral decision-making, while the effects of testosterone are inconsistent. Our study aimed to close this research gap. A sample of 137 cis individuals (73 women, 64 men) completed a moral dilemma task based on the Consequences, Norms, Inaction (CNI) model either in the morning or evening. Relevant personality traits, including agreement with instrumental harm, impartial beneficence, empathy, and gender expression, were assessed via self-report. Blood samples were analyzed for estradiol, progesterone, testosterone, and cortisol. Contrary to prior literature, no behavioral differences between women and men were observed in moral decision-making. However, self-reported agreement with instrumental harm positively predicted sensitivity to consequences and negatively predicted adherence to moral norms in the whole sample. Estradiol concentration positively predicted sensitivity to consequences in the morning women group, and general inaction tendency in the morning men group. Overall, these findings highlight the joint influence of biological and personality factors on moral choices and hint towards subtle sex/gender differences.

## 1. Introduction

Women and men differ in how frequently they choose to use instrumental harm, that is, deliberate infliction of harm as a means to an end, to achieve better outcomes for a greater number of individuals in dilemmatic situations (Friesdorf et al., 2015). One of the common examples of instrumental harm is the Trolley problem: a hypothetical situation, where a trolley is heading towards five individuals on the track but if someone switches the lever, the trolley will change to the track with only one individual on it, resulting in a better overall outcome (Foot, 1978). Moral psychology has long been interested in what principles guide these decisions. Contributions of sex/gender, hormonal profiles, gender expression or moral orientations related to personality to the sex/gender differences in the guiding principles have been investigated previously, however, findings are inconsistent.

Individuals use different heuristics to arrive at a decision in moral dilemmas: one might compare the consequences of each action, or apply commonly accepted moral norms, or one might not engage in moral consideration at all but rather act from their general tendency to engage or disengage in action (Gawronski et al., 2016). The Consequences, Norms, and Generalized Inaction (CNI) model captures these heuristics by employing a factorial design in moral dilemmas. There, either consequences are greater or smaller than costs and moral norms either prohibit or prescribe action. A multinomial modelling is employed to quantify these heuristics into three corresponding behavioral parameters: C parameter for sensitivity to consequences, N parameter for sensitivity to norms, and I parameter for general inaction and action tendency (Gawronski et al., 2017).

The CNI model diverts from the traditional approach of studying moral decision-making. Traditionally, moral behavior used to be operationalized as frequency of choosing instrumental harm in dilemmas where only prohibitive norms are considered, and benefits of an action are always greater if instrumental harm is used. Therefore, using this analytical approach, the inaction and action tendencies were conflated with the moral reasons underlying the choices (Demaree-Cotton & Kahane, 2025; Gawronski et al., 2016).

Sex/gender differences in moral dilemma resolution have been demonstrated first in studies using the traditional approach. Here, men more frequently chose to use instrumental harm than women (Friesdorf et al., 2015). Studies using the CNI model, explicated that women show stronger sensitivity to moral norms and stronger general preference for inaction than men, leading to the difference in frequency of choosing instrumental harm, while men and women do not generally differ in their sensitivity to consequences (Armstrong et al., 2019; Gawronski et al., 2017; Qian et al., 2024).

Gender expression has been suggested to underly the differences in sensitivity to norms and more frequent abstaining from using instrumental harm in women. Higher care orientation, which focuses on prevention of harm to others, has been consistently found in women compared to men across 67 countries (Atari et al., 2020). Care orientation has been also linked to higher empathic concern in women (Dawson et al., 2023) and sex/gender differences in empathic concern have been consistently associated with reduced frequency of choosing instrumental harm (Cordellieri et al., 2020; Fumagalli et al., 2010). However, in Baez et al. (2017), the link between sex/gender differences in choosing instrumental harm and empathic concern depended on self-report measures, suggesting that this link might be affected by the extent by which individuals assume their gender expression in self-reports.

Finally, moral decision-making is altered across several forms of psychopathology as well as acute stress, predisposing individuals either to increased or decreased use of instrumental harm due to altered approach and avoidance tendencies rather than moral inclinations (Ahmadzadeh et al., 2025; Harrison et al., 2012; Koenigs et al., 2012; Yin et al., 2022).

It has also been proposed that different gonadal hormone profiles in women and men contribute to the observed behavioral differences. Prior research in women and men linked testosterone, measured either separately or in relation to cortisol, to increased aggression (Rzepczyk et al., 2024) and reduced empathy (Nitschke & Bartz, 2020; Zilioli et al., 2015). Studies assessing the link between testosterone and the use of instrumental harm have provided inconsistent results, with some studies reporting positive associations between testosterone and instrumental harm (Armbruster et al., 2021; Carney & Mason, 2010; Chen et al., 2016), and others not (Arnocky et al., 2017; Reynolds et al., 2021). Pharmacologically increased testosterone levels in women and men had no effect in most studies using a traditional moral dilemma approach (Arnocky et al., 2017; Chen et al., 2016; Montoya et al., 2013). In Brannon et al. (2019), testosterone administration was positively related to the sensitivity to norms in the CNI model but not to sensitivity to consequences or general inaction/action tendency in both sexes/genders.

The effects of estradiol and progesterone have not been directly assessed before. Yet, these hormones might play a role in moral decision-making as indicated by a study comparing naturally cycling women and oral contraceptive users: here, combined oral contraceptive users more frequently chose to use instrumental harm the longer they used the contraceptives and were more motivated by the consequences to do so (Armbruster et al., 2021). However, it is still not known whether endogenous levels of estradiol or progesterone are related to psychological processes underlying moral decision-making in a similar way.

Estradiol and progesterone influence decision-making behavior through their activational and organizational effects on the human brain (see Ambrase et al., 2021 for review). Importantly, estradiol and progesterone receptor sites are mostly concentrated in brain areas, highly involved in decision-making (Barth et al., 2015). Prior research shows that higher levels of estradiol are positively associated with increased approach behavior and more attention to consequences, while progesterone’s role is less clear (Ambrase et al., 2021). These effects might contribute to sex/gender differences in moral decision-making, especially in modulating women’s sensitivity to consequences and inaction as compared to men.

Prior findings also indicate that cortisol concentration is related to differences in acceptance rates of instrumental harm after acute stress exposure (Kossowska et al., 2016; Singer et al., 2017) and that this association contributes to sex/gender differences in moral decision-making (Singer et al., 2021). Cortisol has a diurnal rhythm, with high concentration in the morning and low in the evening (Czeisler & Klerman, 1999). Importantly, testosterone and estradiol have a similar diurnal fluctuation pattern as cortisol, although the magnitude of these fluctuations differs between women and men (Bao et al., 2003; Brambilla et al., 2009; Grotzinger et al., 2024; Juster et al., 2016). In addition, women exhibit pronounced cyclical fluctuations in estradiol across the menstrual cycle, which exceed the magnitude of the diurnal changes and even alter the timing of the peak level of estradiol (Bao et al., 2003). This suggests that behavioral effects of gonadal hormones might also be dependent on the time of day, while cortisol may represent an important control variable when examining associations between gonadal hormones and moral decision-making.

In this study, we aim to better understand the combined role of women’s and men’s personality and hormonal profiles on their sensitivity to consequences, norms, and inaction tendencies in moral decision-making. We also aim to disentangle any potential effects that might depend on measurement time (morning or evening), as the concentration of the obtained hormones fluctuates during the day. We hypothesize that women will exhibit higher sensitivity to norms and inaction tendency than men. We expect self-reported moral inclination towards the use of instrumental harm to be positively related to sensitivity to consequences both in women and men, while self-reported care orientation and gender expression should be positively associated with the sensitivity to norms in women. We expect to find a positive relationship between endogenous testosterone and sensitivity to consequences in men, and a positive relationship between estradiol and sensitivity to consequences in women. We also expect to find a negative association between estradiol and inaction in women, as well as positive associations between self-reported anxiousness, state anxiety, and behavioral inhibition to inaction in the whole sample.

## 2. Materials and methods

### 2.1. Participants

137 German-native participants between 18 and 35 years of age were recruited from the community. The sample comprised 73 naturally cycling women (M_age_ = 23.04, SD = 2.97) and 64 men (M_age_ = 24.13, SD = 3.73). Advertisements for participation were distributed via mailing lists at the University of Tübingen‘s server as well as through word-of-mouth. Participation was compensated with 15 Euros per hour, two hours maximum. Eligibility criteria included German proficiency at native level.

Individuals with neurological or mental disorders, serious pre-existing medical conditions (e.g., severe hypertension, diabetes, untreated thyroid dysfunction), and current treatment with psychotropic medications, as well as any individuals using hormones (including hormonal contraceptives), who had vasectomy, or who are/were pregnant or breastfeeding within the past year were excluded.

### 2.2. Procedure

The Ethics Commission of the Medical Faculty at the University of Tübingen granted the approval for the study (707/2019BO2, received on 2020-06-17). Potential participants were assessed for inclusion and exclusion criteria via online screening. Participants who successfully met the criteria were invited to the study site (either between 7–8 a.m. or 3–5 p.m.). After arriving at the lab, detailed information about the study was provided, and participants signed an informed consent form and a data protection information sheet. Next, blood was drawn, followed by the assessment of the CNI paradigm presented on a computer screen. Participants provided decisions to moral dilemmas from the CNI paradigm using keys labelled “Yes” and “No” on a standard computer keyboard. Following each dilemma, participants were asked how difficult it was to make the decision in that dilemma (adapted from Conway & Gawronski, 2013). The perceived difficulty was rated using a visual analogue scale from 1 (“not difficult at all”) to 7 (“very difficult”). After completion of the task, participants were also asked “How well were you able to imagine yourself in the dilemmas?”, answering on a scale from 1 (“not well at all”) to 7 (“very well”). After completing the moral dilemmas, participants were asked to complete nine additional psychometric questionnaires reported in section 2.3.2. on a laptop, via the SoSciSurvey platform (www.soscisurvey.de).

### 2.3. Measures

#### 2.3.1. Moral decision-making task

Moral decision-making was assessed with a set of 48 moral dilemmas from the extended Consequences, Norms, and Generalized Inaction Model (CNI model, Körner et al., 2020). The model allows differentiation between various underlying principles for the decision-making (Gawronski et al., 2017; Körner et al., 2020). Within the set of dilemmas, 12 different situations are described, and each situation has four parallel formulations. The formulations differ in whether a proscriptive or a prescriptive norm requires action, and whether the benefits of the action are greater or smaller than the costs. Using multinomial modelling, the CNI model allows for delineating the extent of which the three principles – sensitivity to consequences (C parameter, from 0 to 1, with increasing values corresponding to increasing sensitivity), sensitivity to norms (N parameter, from 0 to 1, with increasing values corresponding to increasing sensitivity) and general inaction and action tendency (I parameter, from 0 to 1, with higher values corresponding to inaction and lower values with action) – underly participant‘s choices.

Two situational scenarios (eight formulations in total) needed to be removed from the analysis due to erroneous recording for all participants, resulting in a total of 10 scenarios (40 variations) assessed.

#### 2.3.2 Self-report measures

##### 2.3.2.1. Gender-related Attributes Survey (GERAS; Gruber et al., 2020)

GERAS is a self-report measure designed to assess individuals’ masculinity and femininity expression. It is comprised of 164 items: 100 personality items expressed as single words, 24 short statements on cognitive ability, and 40 short statements on personal interests and activities, all scored from 1 (not at all) to 7 (very).

##### 2.3.2.2. Moral Foundations Questionnaire (MFQ-30; Graham et al., 2011)

*German translation provided at https://moralfoundations.org/*. Harm/Care orientation was assessed using 6 items depicting compassion and care for others, and disapproval of cruelty and harm. The agreement scores range from 0 to 30, with higher scores indicating higher agreement.

##### 2.3.2.3. Oxford Utilitarianism Scale (OUS; Kahane et al., 2018)

*German translation (Ambrasė et al., 2025)* was used to measure the outcome-based moral deliberation style. The scale consists of two subscales, i.e. Impartial Beneficence (IB-DE, 5 items) and Instrumental Harm (IH-DE, 4 items). The items in the subscales are scored from 1 (strongly disagree) to 7 (strongly agree).

##### 2.3.2.4. Temperament and Character Inventory (TCI; Cloninger et al., 1993)

*German translation (Richter et al., 2000)* was used to measure participants’ inclinations to consider the rewarding consequences of their actions was measured by the reward dependence scale. The scale includes 24 items on social approval, sentimentality, attachment, and dependence. Participants were asked to agree or disagree with the items. The total score for the scale is up to 24 points, and a higher score indicates higher reward dependence.

##### 2.3.2.5. Saarbrücker Personality Questionnaire (SPF; Paulus, 2009)

is a German version of the Interpersonal Reactivity Index (IRI; Davis, 1980). The Empathic Concern subscale was used to assess participants’ empathic concern abilities, because it has been repeatedly related to individual differences in moral decision-making (e.g., Gleichgerrcht & Young, 2013; Körner et al., 2020). The subscale includes 4 items depicting concerned and protective feelings towards others, and uses a 5-point Likert scale, ranging from 1 (does not describe me well) to 5 (describes me very well).

##### 2.3.2.6. Brief Symptom Inventory (BSI; Derogatis, 1993), German version (Franke, 2000)

The Global Severity Index of the BSI was used in this study to assess self-reported current psychological distress level in the participants. The items in the BSI measuring somatization, obsessive-compulsive, interpersonal sensitivity, depression, anxiety, hostility, phobic anxiety, paranoid ideation and psychoticism are scored on a Likert scale from 0 (not at all) to 4 (extremely), indicating the intensity of distress caused by a symptom in the past seven days.

##### 2.3.2.7. Action Regulation Emotion Systems Scale (ARES; Hartig & Moosbrugger, 2003)

ARES measures the Behavioral Inhibition System (BIS) and the Behavioral Approach System (BAS), and is based on the BIS/BAS-Scale (Carver & White, 1994). The BIS scale was used in this study to assess the participants’ proneness to inhibit behavior that might lead to negative outcomes or feelings (sensitivity to punishment). The scale includes two subscales BIS I “Anxiousness” and BIS II “Frustration”. The full scale consists of 23 items which are scored on a Likert scale from 1 (do not agree) to 4 (agree).

##### 2.3.2.8. State-Trait Anxiety Inventory (STAI; Spielberger et al., 1971), German translation (Grimm, 2009)

Current anxious and stressful mood in participants after completing the moral decision-making was measured using the STAI-state subscale. The subscale includes 20 items which are scores from 1 (absolutely not) to 4 (very much).

#### 2.2.3. Hormone assessment

For each participant, 7.5 ml of venous blood was collected in a serum S-Monovette (manufacturer Sarstedt). After blood collection, the sample was sent to the central laboratory of University Hospital Tübingen. There it was centrifuged to obtain plasma, which was aliquoted into microtubes and stored at −70°C. ADVIA Centaur XP assay was used to determine hormone levels of estradiol, progesterone, testosterone, and cortisol (in nmol/L). This assay uses chemiluminescence as the detection method. It shows high precision in the reproducibility of results, with an intra-assay variability of ≤ 7% for cortisol values, ≤ 10% for testosterone, ≤ 5% for estradiol, and ≤ 6% for progesterone values in the range of 0.69–34.70 nmol/L.

### 2.4. Statistical analysis

Descriptive statistics including means and standard deviations, and Pearson correlations were calculated in SPSS Version 28.0.1 (IBM Corporation, 2021). Two participants did not provide blood samples, and two participants did not complete the CNI paradigm; these participants were removed from the analyses where either hormone values or sensitivity to consequences, sensitivity to norms, and general inaction/action tendency were assessed.

For the main analysis, hierarchical regression analyses were conducted in R (version 4.4.2, R Core Team, 2023) to examine the effects of sex/gender, gender expression (femininity and masculinity), time of measurement, hormones (testosterone, estradiol, progesterone, cortisol, all logarithmically transformed due to skewed distribution), and personality factors separately on the C, N, and I parameters as dependent variables. In each model, four-step procedure was applied: 1) Sex/gender, gender expression, and time of measurement (Step 1); 2) Step 1 and hormones; 3) Step 1 and corresponding personality factors; 4) all predictors. The personality factors differed between models based on previous literature (e.g., Atari et al., 2020; Fumagalli et al., 2010). For the sensitivity to consequences (C parameter), OUS Impartial Beneficence, OUS Instrumental Harm, and TCI Reward Dependence were selected as predictors. For the sensitivity to norms (N parameter), we used OUS Impartial Beneficence, OUS Instrumental Harm, MFQ Harm/Care Moral Foundation, and SPF Empathic Concern. For the general inaction/action tendency (I parameter), ARES Behavioral Inhibition, STAI State, and BSI General Severity Index were included. The models were applied to the whole sample, subgroups of all women and men separately, and subgroups of morning and evening sessions for women and men separately. Predictors such as sex/gender, time of measurement, femininity and masculinity, were removed in subgroup analyses where appropriate.

Regression assumptions were evaluated using the Shapiro-Wilk test for normality of residuals, the Breusch-Pagan test for homoscedasticity, and variance inflation factors (VIFs) for multicollinearity. Influential observations were identified using Cook’s distance with a threshold of 0.5. All statistical analyses were performed using R (version 4.3.3.; R Core Team, 2025) and RStudio (version 2026.4.0.526; Posit Team, 2026).

To evaluate the robustness of the regression findings, a sensitivity power analysis using G*Power 3.1.9.7 (Faul et al., 2009) was conducted for those models that revealed significant predictors. The required effect sizes for each sample and each step were calculated assuming α = 0.05, power = 0.8, tested predictor = 1, and appropriate sample sizes and total numbers of predictors. The minimal detectable effect size (f²) for each sample and its step, along with predictors’ partial f^2^ were calculated. If the observed effect size is larger than the minimum detectable effect size (MDES), the findings are considered adequately powered.

Two-way analysis of variance (ANOVA) and independent sample t-tests were used to examine group differences (women vs. men; morning vs. evening women/men) in self-reported dilemma decision difficulty. Decision difficulty was calculated as a mean score (based on Conway & Gawronski, 2013) for the four moral dilemma types included in the CNI model: 1) proscriptive dilemmas where benefits are greater than costs, 2) proscriptive dilemmas where benefits are smaller than costs, 3) prescriptive dilemmas where benefits are greater than costs, 4) prescriptive dilemmas where benefits are smaller than costs.

## 3. Results

### 3.1. Intercorrelations between predictors in the study

Pearson correlation results measuring intercorrelations between the predictors in the study can be found in the supplementary material (Table S2). As expected, sex and testosterone (r = −.997, p < .001) and estradiol (r = .62, p < .001), as well as measurement time and cortisol (r = −.66, p < .001) were strongly intercorrelated. We have dismissed the multicollinearity issue for the regression analyses due to the high intercorrelations between categorical and continuous variables, which could be theoretically anticipated. Other intercorrelations were below the recommended level (r<.6; Yu et al., 2015).

### 3.2. Differences in hormone concentrations

Hormone concentrations (nmol/L), stratified by sex/gender and measurement time, are presented in Table 2 (for convenience of displaying hormone concentrations, Estradiol concentration was converted from pmol/L to nmol/L). Pairwise comparisons for each hormone between groups have been conducted using the Wilcoxon rank-sum test on raw hormonal values.

**Table 1.**
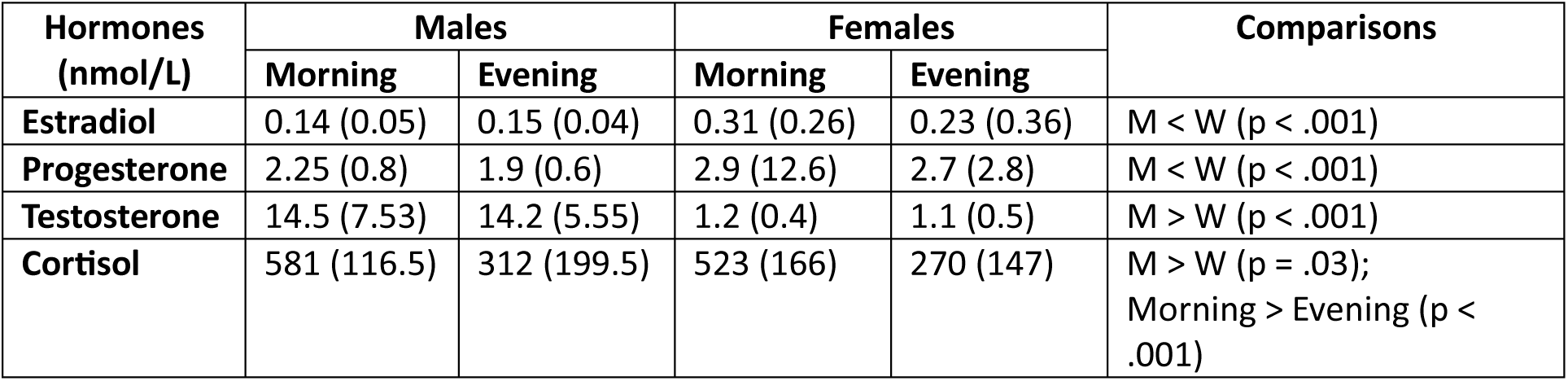
Comparisons of hormonal concentrations stratified by sex and measurement time. Median values with IQR are provided.

**Table 2.** Hierarchical regression results in sensitivity to consequences (C parameter). Effect size for each variable is provided as B (SE). Significant results which survived sensitivity analysis are in bold.

| Predictor | Step 1 | Step 2 | Step 3 | Step 4 |  |  |  |  |  |  |  |  |
| --- | --- | --- | --- | --- | --- | --- | --- | --- | --- | --- | --- | --- |
| Whole sample |  |  |  |  |  |  |  |  |  |  |  |  |
| Sex/gender | -0.002 (0.04) | 0.15 (0.16) | -0.006 (0.04) | 0.16 (0.15) |  |  |  |  |  |  |  |  |
| Time | 0.001 (0.03) | 0.006 (0.05) | 0.003 (0.03) | 0.006 (0.05) |  |  |  |  |  |  |  |  |
| Femininity | -0.03 (0.03) | -0.03 (0.03) | -0.02 (0.03) | -0.02 (0.03) |  |  |  |  |  |  |  |  |
| Masculinity | 0.02 (0.03) | 0.02 (0.03) | 0.02 (0.02) | 0.02 (0.02) |  |  |  |  |  |  |  |  |
| Estradiol |  | 0.01 (0.04) |  | 0.001 (0.04) |  |  |  |  |  |  |  |  |
| Progesterone |  | -0.01 (0.02) |  | 0.004 (0.02) |  |  |  |  |  |  |  |  |
| Testosterone |  | 0.06 (0.06) |  | 0.07 (0.05) |  |  |  |  |  |  |  |  |
| Cortisol |  | 0.002 (0.06) |  | -0.01 (0.05) |  |  |  |  |  |  |  |  |
| OUS Impartial Beneficence |  |  | 0.0004 (0.01) | -0.0002 (0.01) |  |  |  |  |  |  |  |  |
| OUS Instrumental Harm |  |  | **0.06 (0.01)\*\*\*** | **0.07 (0.01)\*\*\*** |  |  |  |  |  |  |  |  |
| TCI Reward dependency |  |  | -0.002 (0.005) | 0.002 (0.006) |  |  |  |  |  |  |  |  |
| R<sup>2</sup> | 0.02 | 0.04 | 0.16 | 0.17 |  |  |  |  |  |  |  |  |
| ΔR<sup>2</sup> |  | 0.02 | 0.12 | 0.01 |  |  |  |  |  |  |  |  |
| F | 0.68 | 0.57 | 3.45 | 2.33 |  |  |  |  |  |  |  |  |
| p | 0.61 | 0.8 | 0.002 | 0.012 |  |  |  |  |  |  |  |  |
|  | Step 1 | Step 2 | Step 3 | Step 4 | Step 1 | Step 2 | Step 3 | Step 4 | Step 1 | Step 2 | Step 3 | Step 4 |
| Entire men group |  |  |  |  | Morning men group |  |  |  | Evening men group |  |  |  |
| Time | 0.009 (0.05) | -0.003 (0.07) | 0.014 (0.05) | -0.02 (0.07) |  |  |  |  |  |  |  |  |
| Masculinity | 0.03 (0.04) | 0.03 (0.04) | 0.03 (0.04) | 0.03 (0.04) | 0.06 (0.05) | 0.05 (0.06) | 0.06 (0.05) | 0.05 (0.05) | 0.01 (0.05) | 0.01 (0.06) | -0.01 (0.05) | -0.03 (0.06) |
| Estradiol |  | 0.02 (0.13) |  | -0.14 (0.12) |  | -0.06 (0.18) |  | -0.09 (0.17) |  | 0.16 (0.23) |  | 0.02 (0.21) |
| Progesterone |  | 0.08 (0.11) |  | 0.12 (0.1) |  | 0.12 (0.16) |  | 0.12 (0.14) |  | -0.01 (0.17) |  | 0.12 (0.16) |
| Testosterone |  | 0.06 (0.09) |  | 0.17 (0.09) |  | 0.12 (0.14) |  | 0.06 (0.13) |  | -0.02 (0.15) |  | 0.22 (0.15) |
| Cortisol |  | -0.05 (0.1) |  | -0.12 (0.09) |  | 0.09 (0.27) |  | 0.16 (0.22) |  | -0.06 (0.12) |  | -0.11 (0.11) |
| OUS Impartial Beneficence |  |  | 0.002 (0.02) | 0.004 (0.02) |  |  | 0.06 (0.03)* | 0.06 (0.03)* |  |  | -0.05 (0.03) | -0.04 (0.03) |
| OUS Instrumental Harm |  |  | **0.07 (0.02)\*\*** | **0.08 (0.02)\*\*\*** |  |  | **0.09 (0.03)\*\*** | **0.09 (0.03)\*\*** |  |  | 0.04 (0.03) | 0.06 (0.03) |
| TCI Reward dependency |  |  | -0.003 (0.007) | -0.006 (0.008) |  |  | 0.009 (0.009) | 0.009 (0.01) |  |  | -0.01 (0.01) | -0.02 (0.01) |
| R<sup>2</sup> | 0.01 | 0.03 | 0.2 | 0.27 | 0.04 | 0.09 | 0.4 | 0.46 | 0.001 | 0.04 | 0.3 | 0.41 |
| ΔR<sup>2</sup> |  | 0.02 | 0.17 | 0.07 |  | 0.05 | 0.31 | 0.06 |  | 0.039 | 0.26 | 0.11 |
| F | 0.45 | 0.31 | 2.89 | 2.14 | 1.24 | 0.52 | 4.39 | 2.43 | 0.03 | 0.19 | 2.89 | 1.88 |
| p | 0.64 | 0.93 | 0.02 | 0.042 | 0.27 | 0.76 | 0.007 | 0.046 | 0.86 | 0.96 | 0.04 | 0.11 |
| Entire women group |  |  |  |  | Morning women group |  |  |  | Evening women group |  |  |  |
| Time | -0.005 (0.07) | -0.0001 (0.07) | -0.007 (0.04) | 0.02 (0.07) |  |  |  |  |  |  |  |  |
| Femininity | -0.02 (0.04) | -0.02 (0.04) | -0.03 (0.04) | -0.03 (0.04) | -0.11 (0.06) | -0.13 (0.07) | -0.09 (0.06) | -0.09 (0.06) | 0.02 (0.05) | 0.02 (0.05) | 0.007 (0.05) | -0.001 (0.06) |
| Estradiol |  | 0.02 (0.04) |  | 0.02 (0.04) |  | 0.15 (0.06)* |  | **0.18 (0.06)\*\*** |  | -0.08 (0.06) |  | -0.07 (0.06) |
| Progesterone |  | -0.01 (0.03) |  | 0.0004 (0.03) |  | -0.08 (0.03)* |  | -0.07 (0.03) |  | 0.02 (0.04) |  | 0.03 (0.04) |
| Testosterone |  | 0.06 (0.08) |  | 0.04 (0.08) |  | 0.02 (0.11) |  | 0.07 (0.1) |  | 0.14 (0.11) |  | 0.07 (0.11) |
| Cortisol |  | 0.003 (0.08) |  | 0.03 (0.07) |  | -0.04 (0.11) |  | -0.04 (0.11) |  | -0.01 (0.09) |  | 0.05 (0.09) |
| OUS Impartial Beneficence |  |  | -0.0004 (0.02) | -0.004 (0.02) |  |  | 0.02 (0.03) | 0.02 (0.03) |  |  | -0.03 (0.04) | -0.02 (0.04) |
| OUS Instrumental Harm |  |  | **0.06 (0.02)\*\*** | **0.06 (0.02)\*** |  |  | 0.05 (0.03) | 0.06 (0.03) |  |  | 0.06 (0.03) | 0.06 (0.03) |
| TCI Reward dependency |  |  | 0.006 (0.007) | 0.007 (0.008) |  |  | 0.001 (0.01) | 0.004 (0.01) |  |  | 0.008 (0.01) | 0.01 (0.01) |
| R<sup>2</sup> | 0.006 | 0.02 | 0.12 | 0.14 | 0.09 | 0.32 | 0.21 | 0.47 | 0.004 | 0.10 | 0.13 | 0.2 |
| ΔR<sup>2</sup> |  | 0.014 | 0.1 | 0.02 |  | 0.23 | -0.11 | 0.26 |  | 0.096 | 0.03 | 0.07 |
| F | 0.19 | 0.24 | 1.77 | 1.04 | 3.28 | 2.56 | 1.96 | 2.65 | 0.15 | 0.66 | 1.19 | 0.88 |
| p | 0.83 | 0.96 | 0.13 | 0.42 | 0.08 | 0.05 | 0.13 | 0.03 | 0.70 | 0.66 | 0.33 | 0.54 |

### 3.3. Sex/gender differences in the CNI model

Two-way ANOVA results suggest that there are no between-subjects effects of sex/gender or measurement time or their interaction in sensitivity to consequences (C parameter) in our sample (all Fs < .92, ps > .34, see Supplemental Tables S3 and S4).

For sensitivity to norms (N parameter), we found a significant effect of measurement time (F(1,131) = 6.5, p = .01, 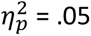, indicating that participants measured in the morning, were more sensitive to moral norms than participants measured in the evening. No sex/gender effect or interaction of measurement time * sex/gender emerged (all Fs < 1.92, ps > .17, see Supplemental Tables S5 and S6).

Regarding the I parameter, no between-subjects effects of sex/gender or measurement time or their interaction were found for general inaction/action tendency (all Fs < .95, ps > .33, see Supplemental Tables S7 and S8).

### 3.4. Hierarchical regression analyses

#### 3.4.1. Prediction of the sensitivity to consequences – C parameter

Hierarchical multiple regressions were conducted to examine predictors of sensitivity to consequences (C parameter) as dependent variable. Personality factors in these analyses included OUS Impartial Beneficence, OUS Instrumental Harm, and TCI Reward Dependence. Diagnostics of linear regressions did not indicate substantial violations of model assumptions (see Supplemental Table S8).

Table 3 presents the statistical details from the hierarchical linear regression analyses. In the whole sample we found statistically significant positive associations between sensitivity to consequences (C parameter) and OUS Instrumental Harm consistently (Step 4: B = 0.07, SE = 0.01, t = 4.32, p < .001), and in women and men separately. No effects of sex/gender, gender expression (femininity and masculinity), measurement time, and hormone levels were identified in these groups (all ps > .053).

**Table 3.** Hierarchical regression results in sensitivity to norms (N parameter). Effect size for each variable is provided as B (SE). Significant results which survived sensitivity analysis are in bold.

| Predictor | Step 1 | Step 2 | Step 3 | Step 4 |
| --- | --- | --- | --- | --- |
| **Whole sample** |  |  |  |  |
| Sex | 0.03 (0.06) | 0.09 (0.26) | 0.008 (0.06) | 0.06 (0.25) |
| Time | -0.14 (0.05)* | -0.12 (0.08) | -0.14 (0.05)** | -0.1 (0.07) |
| Femininity | 0.02 (0.04) | 0.01 (0.05) | -0.02 (0.05) | -0.04 (0.05) |
| Masculinity | 0.001 (0.04) | 0.001 (0.04) | 0.007 (0.04) | 0.006 (0.04) |
| Estradiol |  | 0.08 (0.07) |  | 0.08 (0.06) |
| Progesterone |  | -0.04 (0.04) |  | -0.06 (0.04) |
| Testosterone |  | 0.04 (0.09) |  | 0.03 (0.09) |
| Cortisol |  | 0.03 (0.09) |  | 0.07 (0.09) |
| Impartial Beneficence |  |  | 0.02 (0.02) | 0.01 (0.03) |
| Instrumental Harm |  |  | **-0.07 (0.02)\*\*** | -0.08 (0.02)** |
| Care/Harm foundation |  |  | 0.04 (0.04) | 0.02 (0.04) |
| Empathic concern |  |  | 0.01 (0.01) | 0.02 (0.01) |
| **R<sup>2</sup>** | 0.05 | 0.07 | 0.16 | 0.18 |
| **ΔR<sup>2</sup>** |  | 0.02 | 0.09 | 0.02 |
| **F** | 1.86 | 1.08 | 3.01 | 2.23 |
| **p** | 0.12 | 0.38 | 0.004 | 0.01 |

|  | Step 1 | Step 2 | Step 3 | Step 4 | Step 1 | Step 2 | Step 3 | Step 4 | Step 1 | Step 2 | Step 3 | Step 4 |
| --- | --- | --- | --- | --- | --- | --- | --- | --- | --- | --- | --- | --- |
| **Entire men group** |  |  |  |  | **Morning men group** |  |  |  | **Evening men group** |  |  |  |
| Time | -0.06 (0.08) | -0.02 (0.12) | -0.04 (0.08) | -0.006 (0.12) |  |  |  |  |  |  |  |  |
| Masculinity | -0.004 (0.06) | -0.007 (0.06) | 0.006 (0.06) | 0.003 (0.06) | 0.04 (0.08) | 0.07 (0.08) | -0.03 (0.08) | -0.03 (0.09) | -0.05 (0.10) | -0.04 (0.11) | -0.09 (0.10) | -0.08 (0.11) |
| Estradiol |  | -0.12 (0.21) |  | -0.04 (0.22) |  | -0.05 (0.23) |  | 0.27 (0.24) |  | -0.38 (0.39) |  | -0.10 (0.49) |
| Progesterone |  | -0.17 (0.18) |  | -0.21 (0.17) |  | -0.41 (0.21) |  | -0.18 (0.21) |  | 0.14 (0.29) |  | 0.009 (0.32) |
| Testosterone |  | 0.02 (0.15) |  | -0.08 (0.15) |  | 0.23 (0.19) |  | 0.12 (0.18) |  | 0.006 (0.25) |  | -0.11 (0.27) |
| Cortisol |  | 0.07 (0.16) |  | 0.1 (0.16) |  | -0.17 (0.35) |  | -0.31 (0.33) |  | 0.03 (0.20) |  | 0.14 (0.23) |
| Impartial Beneficence |  |  | 0.03 (0.04) | 0.02 (0.04) |  |  | -0.02 (0.05) | -0.04 (0.05) |  |  | 0.03 (0.05) | 0.03 (0.06) |
| Instrumental Harm |  |  | -0.07 (0.03) | -0.07 (0.04) |  |  | -0.09 (0.04)* | -0.10 (0.04)* |  |  | -0.11 (0.06) | -0.13 (0.08) |
| Care/Harm foundation |  |  | 0.02 (0.06) | 0.02 (0.06) |  |  | 0.19 (0.08) * | 0.19 (0.09)* |  |  | -0.16 (0.10) | -0.15 (0.12) |
| Empathic concern |  |  | 0.02 (0.01) | 0.02 (0.02) |  |  | 0.002 (0.02) | -0.008 (0.02) |  |  | 0.02 (0.02) | 0.02 (0.02) |
| **R<sup>2</sup>** | 0.01 | 0.03 | 0.13 | 0.16 | 0.007 | 0.25 | 0.32 | 0.46 | 0.008 | 0.08 | 0.19 | 0.25 |
| **ΔR<sup>2</sup>** |  | 0.02 | 0.1 | 0.03 |  | 0.243 | 0.07 | 0.14 |  | 0.072 | 0.11 | 0.06 |
| **F** | 0.29 | 0.28 | 1.41 | 0.99 | 0.22 | 1.69 | 2.40 | 2.12 | 0.24 | 0.42 | 1.22 | 0.79 |
| **p** | 0.75 | 0.94 | 0.23 | 0.47 | 0.64 | 0.17 | 0.06 | 0.07 | 0.63 | 0.83 | 0.33 | 0.63 |
| **Entire women group** |  |  |  |  | **Morning women group** |  |  |  | **Evening women group** |  |  |  |
| Time | **-0.21 (0.07)\*\*** | -0.19 (0.1) | **-0.25 (0.07)\*\*** | -0.2 (0.11) |  |  |  |  |  |  |  |  |
| Femininity | 0.02 (0.06) | 0.02 (0.06) | 0.01 (0.06) | -0.02 (0.07) | 0.07 (0.09) | 0.005 (0.11) | 0.10 (0.10) | -0.04 (0.12) | 0.0007 (0.08) | 0.002 (0.08) | -0.07 (0.09) | -0.06 (0.09) |
| Estradiol |  | 0.1 (0.07) |  | 0.08 (0.07) |  | 0.04 (0.10) |  | 0.02 (0.10) |  | 0.14 (0.10) |  | 0.08 (0.12) |
| Progesterone |  | -0.05 (0.04) |  | -0.07 (0.04) |  | -0.07 (0.05) |  | -0.10 (0.05) |  | 0.004 (0.06) |  | 0.004 (0.07) |
| Testosterone |  | 0.07 (0.12) |  | 0.12 (0.12) |  | 0.08 (0.17) |  | -0.01 (0.17) |  | -0.005 (0.17) |  | 0.10 (0.18) |
| Cortisol |  | 0.03 (0.11) |  | 0.05 (0.12) |  | 0.20 (0.18) |  | 0.22 (0.19) |  | -0.02 (0.14) |  | 0.0003 (0.17) |
| Impartial Beneficence |  |  | 0.04 (0.04) | 0.03 (0.04) |  |  | 0.01 (0.05) | 0.003 (0.05) |  |  | 0.09 (0.06) | 0.08 (0.08) |
| Instrumental Harm |  |  | -0.07 (0.04) | -0.08 (0.04)* |  |  | -0.06 (0.05) | -0.11 (0.06) |  |  | -0.06 (0.05) | -0.08 (0.05) |
| Care/Harm foundation |  |  | 0.08 (0.06) | 0.05 (0.07) |  |  | 0.09 (0.09) | 0.06 (0.10) |  |  | 0.07 (0.10) | 0.03 (0.11) |
| Empathic concern |  |  | -0.0003 (0.01) | 0.01 (0.02) |  |  | -0.02 (0.02) | 0.0002 (0.02) |  |  | 0.02 (0.02) | 0.02 (0.03) |
| **R<sup>2</sup>** | 0.12 | 0.16 | 0.24 | 0.28 | 0.02 | 0.15 | 0.14 | 0.33 | 0.000002 | 0.11 | 0.23 | 0.29 |
| **ΔR<sup>2</sup>** |  | 0.04 | 0.08 | 0.04 |  | 0.13 | -0.01 | 0.19 |  | 0.109998 | 0.12 | 0.06 |
| **F** | 4.47 | 1.97 | 3.27 | 2.27 | 0.52 | 0.96 | 0.92 | 1.24 | 0.00007 | 0.74 | 1.81 | 1.16 |
| **p** | 0.02 | 0.08 | 0.007 | 0.03 | 0.48 | 0.46 | 0.48 | 0.32 | 0.99 | 0.60 | 0.14 | 0.36 |

When the sample was split by sex/gender and measurement time, different predictors emerged in each group: in the morning men group, OUS Impartial Beneficence and OUS Instrumental Harm positively predicted sensitivity to consequences at Step 3 and Step 4 (Step 3: R² = .39, R²_adj_ = .3, F(4, 27) = 4.39, p = .007; OUS Instrumental Harm: B = 0.09, SE = 0.03, t = 3.28, p = .003, OUS Impartial Beneficence: B = 0.06, SE = 0.03, p = 0.03). No significant predictors emerged in the evening men group (all ps > .058).

In the morning women group, estradiol significantly positively and progesterone significantly negatively predicted sensitivity to consequences (Step 2: R² = .32, R²_adj_ = .2, F(5, 27) = 2.56, p = .05; estradiol: B = 0.15, SE = 0.06, t = 2.46, p = .02; progesterone: B = −0.08, SE = 0.03, t = −2.58, p = .02). Association between estradiol and sensitivity to consequences remained significant in the last step of the model when hormones and personality factors were included (Step 4: R² = .47, R²_adj_ = .29, F(8, 24) = 2.65, p = .03; B = 0.18, SE = 0.06, t = 3.0, p = .006). Here, model fit was improved, and the model remained significant. No significant predictors emerged for the evening women group (all ps > .08).

According to the sensitivity analysis results (Supplemental Table S12), the effect sizes for OUS Instrumental Harm in the whole sample, the entire men group, and the morning men group, along with the effect size for estradiol in the full model of the morning women group, exceeded the minimal detection threshold. The effect sizes of the other predictors (OUS Impartial Beneficence and progesterone) did not exceed the detection threshold.

Taken together, our analyses demonstrated that across the full sample, individuals who showed greater agreement with OUS Instrumental Harm also tended to show greater sensitivity to consequences (C parameter). This relationship remained significant after considering sex/gender, gender expression (femininity and masculinity), and hormone concentrations. Among women tested in the morning, higher estradiol concentrations were linked to greater sensitivity to consequences, independent of the other variables included in the model.

#### 3.4.2. Prediction of the sensitivity to norms

Hierarchical multiple regressions were conducted to examine predictors of sensitivity to norms (N parameter) as dependent variable. Personality factors in these analyses included OUS Impartial Beneficence, OUS Instrumental Harm, MFQ Harm/Care Foundation, and SPF Empathic Concern. Diagnostics of linear regressions did not indicate substantial violations of model assumptions (see Supplemental Table S9).

Table 4 presents the statistical details from the hierarchical linear regression analyses. We found a significant negative association between sensitivity to norms and OUS Instrumental Harm as well as measurement time in the whole sample at Step 3 (R² = .16, R²_adj_ = .11, F(8,125) = 3.01, p = .004; Time: B = −0.14, SE = 0.05, t = −2.67, p = .009, OUS Instrumental Harm: B = −0.07, SE = 0.02, t = −3.06, p = .003).

**Table 4.**
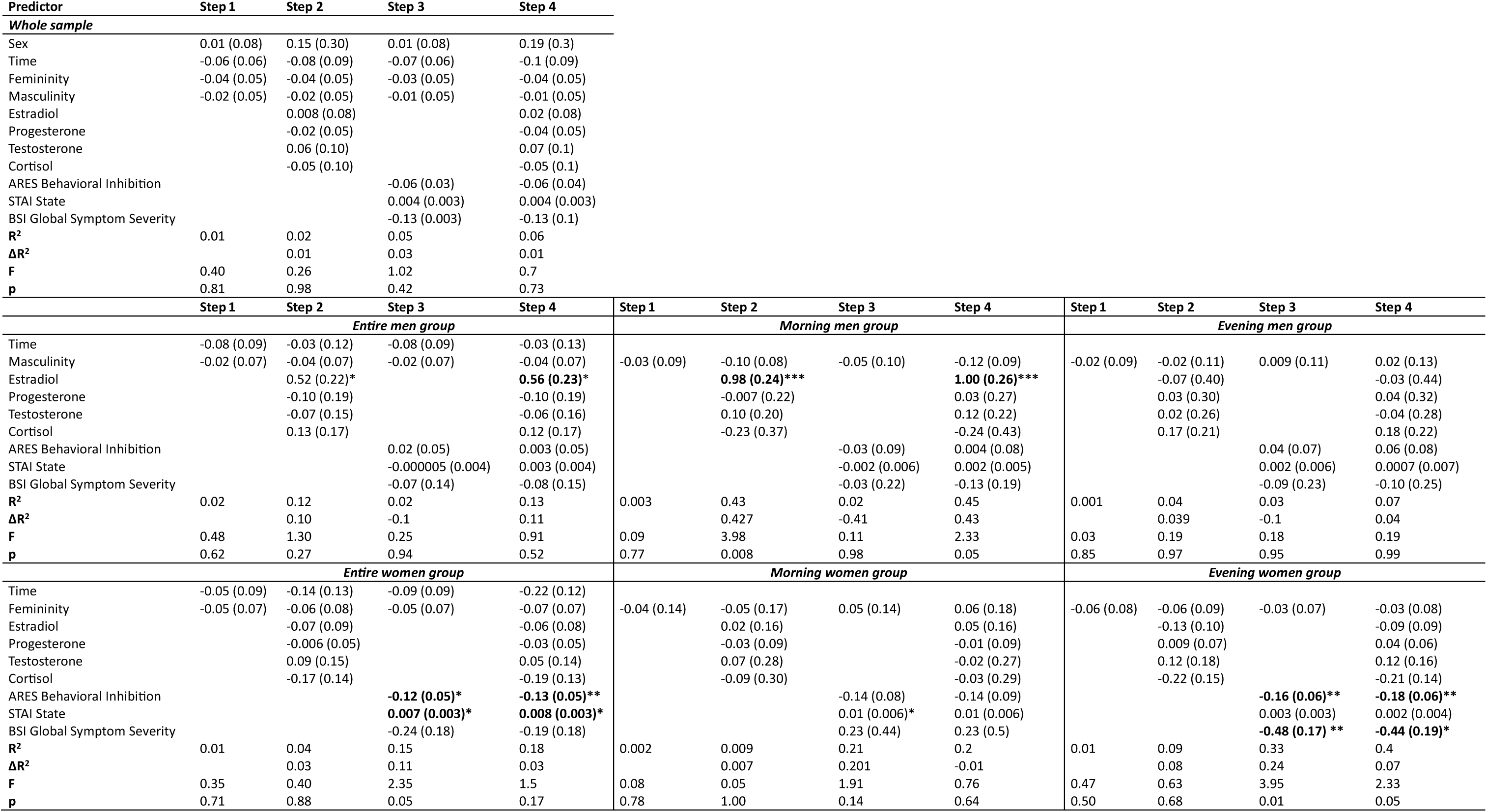
Hierarchical regression results in general inaction/action tendency (I parameter). Effect size for each variable is provided as B (SE). Significant results which survived sensitivity analysis are in bold.

For men, OUS Instrumental Harm together with MFQ Harm/Care foundation emerged as significant predictors of sensitivity to norms only in the morning group at Step 3 (R² = .32, R²_adj_ = .18, F(5,26) = 2.70, p = .06; OUS Instrumental Harm: B = −0.09, SE = 0.04, t = −2.22, p = .004), MFQ Care/Harm foundation: B = 0.19, SE = 0.08, t = 2.30, p = .003), and they remained significant when hormones were included in the model (Step 4: R² = .46, R²_adj_ = .25, F(9,22) = 2.12, p = .07; OUS Instrumental Harm: B = −0.10, SE = 0.04, t = −2.39, p = .003; MFQ Care/Harm Foundation: B = 0.19, SE = 0.09, t = 2.21, p = .004). However, the model remained insignificant in both steps. No significant associations were found for the entire men group and the evening men subgroup.

In the entire women’s group, OUS Instrumental Harm remained a significant predictor but only in combination with the hormones at Step 4 (R² = .28, F(10,58) = 2.27, p = .03; B = −0.08, SE = 0.04, t = −2.22, p = .03). Here, measurement time also emerged as a significant predictor in the steps where personality factors were not included (Step 1: R² = .12, R²_adj_ = .09, F(2,68) = 4.47, p = .02; B = −0.21, SE = 0.07, t = −2.92, p = .005; Step 3: R² = .24, R²_adj_ = .16, F(6,63) = 3.27, p = .007; B = −0.25, SE = 0.07, t = −3.39, p = .001), indicating that there might be a group difference between women measured in the morning and in the evening. Subsequent analyses in these subgroups did not find any significant predictors (all ps > .07).

According to the sensitivity analysis results (Supplemental Table S12), only the effect size for OUS Instrumental Harm in Step 3 of the whole sample and effect sizes for measurement time in Steps 1 and 3 of the entire women sample exceeded the minimal detection threshold. The effect sizes of the other predictors did not exceed the detection threshold.

Taken together, greater agreement with OUS Instrumental Harm was associated with lower sensitivity to norms across the full sample, and participants tested in the evening showed lower sensitivity to norms than those tested in the morning. In the women’s sample, OUS Instrumental Harm remained a significant negative predictor of sensitivity to norms when hormone concentrations were included in the model, while measurement time was associated with sensitivity to norms across several models when hormone concentrations were not accounted for. Although OUS Instrumental Harm and the MFQ Care/Harm foundation were associated with sensitivity to norms among men tested in the morning, these findings should be interpreted cautiously as the overall models were not statistically significant, and sensitivity analyses suggested that only the effects of Instrumental Harm in the full sample and measurement time in the women’s sample were sufficiently robust.

#### 3.4.3. Prediction of the general inaction/action tendency

Hierarchical multiple regressions were conducted to examine predictors of sensitivity to general inaction/action tendency (I parameter) as dependent variable. Personality factors in these analyses included ARES Behavioral Inhibition, STAI State, and BSI Global Symptom Severity. Diagnostics of linear regressions indicated influential data points in the morning men’s group (see Supplemental Table S11).

Table 5 presents the statistical details from the hierarchical linear regression analyses. In our analysis of the whole sample, we did not find any significant predictors for the general inaction/action tendency (all ps > .07). However, different significant predictors emerged for women and men separately. In the entire men group, estradiol had a moderate positive association with general inaction/action tendency (Step 4: R² = .13, R²_adj_ = −.01, F(9, 53) = 0.91, p = .52; B = 0.56, SE = 0.23, t = 2.45, p = .02), yet the model did not reach statistical significance. Splitting the group further according to measurement time indicated that this result was driven by the morning men subgroup (Step 2: R² = .43, R²_adj_ = .32, F(5,26) = 3.98, p = .008; B = 0.80, SE = 0.24, t = 4.06, p = .0004). Adding personality factors decreased the model fit (Step 4: R² = .45, R²_adj_ = .25, F(8, 23) = 2.33, p = .054; B = 1.0, SE = 0.26, t = 3.87, p = .0008), indicating that the prediction step including masculinity and hormones should be favored.

In the entire women group, ARES Behavioral Inhibition negatively and STAI State positively predicted the general inaction/action tendency at Step 3 (R² = .15, R²_adj_ = .09, F(5, 65) = 2.40, p = .05; ARES Behavioral Inhibition: B = −0.12, SE = 0.05, t = −2.53, p = .01; STAI State: B = 0.007, SE = 0.003, t = 2.24, p = .03). These predictions remained significant also when the hormones were added, yet at this step, the model became insignificant (Step 4: R² = .18, R²_adj_ = .06, F(9, 60) = 1.5, p = .17). After further splitting the group by measurement time, we found that STAI State remained a significant predictor in the morning women subgroup (Step 3: R² = .21, R²_adj_ = .13, F(4,29) = 2.23, p = .1; B = 0.01, SE = 0.006, t = 2.19, p = .04), yet the model was not significant. In the evening women group, contrary to our expectations, ARES Behavioral Inhibition and BSI General Symptom Severity were significantly and negatively associated with the general inaction/action tendency at Step 3 (R² = .33, R²_adj_ = .25, F(4, 32) = 3.95, p = .01; ARES Behavioral Inhibition: B = −0.16, SE = 0.06, t = −2.89, p = .007, BSI General Symptom Severity: B = −0.48, SE = 0.17, t = −2.79, p = .009). Controlling for the hormones along the personality factors decreased the model fit (Step 4: R² = .4, R²_adj_ = .23, F(8, 28) = 2.33, p = .046) but the two personality predictors from the previous step remained significant (ARES Behavioral Inhibition: B = −0.18, SE = 0.06, t = −3.07, p = .005, BSI General Symptom Severity: B = −0.44, SE = 0.19, t = −2.36, p = .03).

According to the sensitivity analysis results (Supplemental Table S12), the effect sizes for estradiol in the morning men group and for ARES Behavioral Inhibition in the entire women group, as well as in the evening women group together with BIS General Symptom Severity, exceeded the minimal detection threshold. The effect sizes of the other predictors did not exceed the detection threshold.

Taken together, no significant predictors of general inaction/action tendency (I parameter) emerged in the full sample. However, analyses within sex/gender-specific subgroups suggested different reliable patterns: among men, higher estradiol concentrations were associated with a greater tendency toward inaction, particularly in the morning subgroup, whereas among women, higher ARES Behavioral Inhibition and BIS General Symptom Severity were associated with a lower tendency toward inaction.

### 3.5. Group differences in dilemma decision difficulty

After each dilemma, we also assessed difficulty in making decisions in women and men. We were particularly interested whether group or time effects emerge for the four categories of dilemma scenarios in the CNI model.

Running mixed ANOVAs, we found sex/gender effects on decision difficulty in incongruent scenarios where proscriptive norms prohibited harmful action and benefits of that action were greater than costs (F(1, 132) = 4.76, p = .03, *η*^2^ = .04). Here, women found it more difficult than men to arrive at the decision (see Supplemental Tables S13 and S14).

Sex/gender also affected how difficult participants reported to make decisions in incongruent scenarios where prescriptive norms prescribed prevention of harm, yet its benefits were smaller than costs (F(1, 132) = 5.18, p = .02, *η*^2^ = .04). Again, women reported having more difficulty to make decisions than men (see Supplemental Tables S18 and S19). We did not find any further main or interaction effects (all ps > .16, see Supplemental Tables S15 - S18).

## 4. Discussion

In this study, we wanted to better understand the combined role of women’s and men’s personality and hormonal profiles in moral decision-making, specifically their sensitivity to consequences and norms, and inaction tendencies. We did not find associations between sex/gender or gender expression and moral decision-making heuristics. However, we observed first evidence that estradiol is associated with moral decision-making, with differential effects in women and men and time of measurement. Additionally, personality factors were associated with moral reasoning in women and men.

### 4.1. No differences in moral heuristics between women and men

Based on previous literature (Armstrong et al., 2019; Gawronski et al., 2017; Qian et al., 2024), we tested whether women and men differ in how they reason during moral decision-making using the CNI model. It allows researchers to disentangle psychological processes – sensitivity to consequences, sensitivity to norms, or general tendency towards inaction/action – underlying participants’ moral choices by utilizing multinomial modelling. Our results showed no difference between women and men when we directly compared the groups in any of the CNI model parameters. Furthermore, we also included their self-reported femininity and masculinity expression in the hierarchical regression models but again did not observe significant predictions for any of the CNI model parameters.

Our results divert from the findings of moderate-to-strong effects of sex/gender in previous studies, where women exhibited higher sensitivity to norms and higher general inaction/action tendency in the CNI model than men (Gawronski et al., 2017). Several factors may contribute to this divergence and should be considered in further investigations of sex/gender differences: influence of the region where the measurement takes place, education, the mode of measurement, i.e., online or in person, and the age of participants. In the study by Gawronski et al. (2017), somewhat older (M_age_ = 32.2 years, SD = 10.96) English-native speakers were recruited for the study via MTurk survey platform and performed it online. In our study participants on average were younger (M_age_ = 23.6 years, SD = 3.38) German-native students with possibly a different educational background and performed the experiment at the laboratory where an examiner was present in the same room.

Cultural differences, age, and presence of other individuals during moral decision-making have been shown to influence moral decision-making (Atari et al., 2020; Awad et al., 2020; FeldmanHall et al., 2012; Lin et al., 2024; Pilcher & Smith, 2024). Importantly, the presence of another individual during the experiment affected men more and decreased their use of instrumental harm in moral dilemmas (Pilcher & Smith, 2024). While we cannot conclude that our experimental conditions impacted women’s and men’s moral choices, future studies should take this into account and systematically investigate this factor.

We did, however, find that women had more difficulty making decisions in incongruent moral scenarios, that is, scenarios in which moral norms and consequences conflicted. Specifically, women found it more difficult than men to decide when moral norms prohibited instrumental harm despite the action yielding greater overall benefits, and when moral norms prescribed preventing harm despite the action yielding smaller benefits than costs.

Our findings are in line with other studies using different moral decision-making paradigms. Previously, a meta-analysis in a large sample of respondents (N = 1,837) has demonstrated that women rated incongruent moral dilemmas as more difficult than men, yet the effect size of this difference was small (d = .38, SE = .05; Friesdorf et al., 2015). Similarly, women reported greater emotional distress when accepting instrumental harm options in incongruent dilemmas (Cordellieri et al., 2020). This mirrors results of earlier studies where women were more hesitant to use instrumental harm for greater benefit in more emotionally salient moral dilemmas only (Capraro & Sippel, 2017; Fumagalli et al., 2010). Based on these previous findings and our results, one could speculate that the reported differences in difficulty might be contributing to sex/gender differences in moral choices in incongruent scenarios only and lesser so to psychological mechanisms underlying those choices.

### 4.2. Sex/gender-specific effect of estradiol in moral heuristics

In this study, we found that estradiol affects moral heuristics during decision-making in a sex/gender-specific manner. Higher estradiol was related to higher sensitivity to consequences in women and a stronger tendency toward inaction in men, both groups measured in the morning. To our knowledge, this is the first study to examine associations between endogenous estradiol concentration and moral decision-making heuristics in both women and men, thereby extending current understanding of the hormonal correlates of moral judgment.

#### 4.2.1. Estradiol effect in women

In this study, higher estradiol was associated with higher sensitivity to consequences in the morning women group. Only one previous study investigated estradiol and progesterone effects on moral decision-making, by comparing naturally cycling women (early follicular or late luteal cycle phase), women who use oral contraceptives, and men (Armbruster et al., 2021). They found that women using combined oral contraceptives more frequently agreed to use instrumental harm than naturally cycling women but less frequently than men. Combined oral contraceptives usually contain synthetic ethinyl estradiol and synthetic progestin. Ethinyl estradiol as compared to endogenous Estradiol (17β-estradiol) has a stronger receptor affinity and biological potency (Stanczyk et al., 2024). Synthetic progestins might have a lower or higher receptor affinity and they activate progesterone and androgen receptors in the brain (García-Sáenz et al., 2023). Through these mechanisms, combined oral contraceptives alter functional activation of brain regions responsible for emotion, mood and cognition (Kimmig et al., 2023; Lewis et al., 2019). These effects might contribute to the behavioral difference in oral contraceptive users and naturally cycling women in the previously cited study. Notably, despite differences in hormonal milieu, the direction of the association observed in our study is consistent with the findings by Armbruster et al. (2021), which suggest that higher estradiol concentrations may be associated with greater acceptance of instrumental harm through increased sensitivity to consequences.

This interpretation is also supported by findings from other value-based decision-making domains. Estradiol has repeatedly been implicated in valuation and reward-processing systems. Women are more likely to choose high-reward options despite elevated risk (Lazzaro et al., 2016), demonstrate improved reward maximization by allocating greater effort to optimize outcomes (Jakob et al., 2018), and increased their effort to obtain rewards faster (Grahlow et al., 2026) during late follicular phase, when estradiol concentrations peak. Furthermore, avoidance learning decreases during the follicular phase as estradiol levels rise, suggesting reduced sensitivity to potential negative outcomes and greater attention to potential rewards or benefits even at the possibility of negative outcomes (Diekhof et al., 2020). However, as Grahlow et al. (2026) emphasizes, in women, associations between estradiol and changes in behavior might not depend on the absolute levels of the hormone but rather on the menstrual cycle phase. These effects were not assessed in our study.

Taken together, these findings suggest that estradiol might shape decision-making not simply by altering preferences for risk or reward, but by changing how strongly consequences are represented, valued, and integrated into action selection. Within the context of moral decision-making, such mechanisms could manifest as greater consideration of outcomes when evaluating whether instrumental harm is justified, which is consistent with the positive association between estradiol and consequence sensitivity observed in the present study.

#### 4.2.2. Estradiol effect in men

We also found that higher estradiol was associated with higher inaction tendency in the morning men group. The behavioral functions of estradiol in men’s decision-making have been very rarely studied, and our study related estradiol to individual differences regarding inaction and action tendencies in moral decision-making in men for the first time.

Contrary to our results, estradiol has been associated with higher moral sensitivity regarding economic fairness in men (Coenjaerts et al., 2021). In decision-making, its effects paralleled the direction of estradiol effects in women by increasing reward sensitivity and salience of previous choice outcomes in men (Veselic et al., 2021). It has been also related to higher use of harm when reacting aggressively to provocation (for a meta-analysis, see Wang et al., 2023).

Our finding is, however, broadly consistent with previous research linking higher estradiol in males to less externally directed behavior. For example, estradiol has been associated with reduced violent physical alcohol-related aggression in adult men during morning measurements (Eriksson et al., 2003), lower development of aggressive behavior in youth with higher baseline estradiol levels (Azurmendi et al., 2016), and lower sensation seeking during adolescence and young adulthood (Harden et al., 2018).

Thus, these findings raise the possibility that higher estradiol is associated with behavioral tendencies favoring restraint over action in some specific contexts, although further research is needed to determine whether the mechanisms underlying reduced aggression and sensation seeking are related to the increased inaction tendencies observed in the present study.

#### 4.2.3. Diurnal estradiol effects

An important aspect of our estradiol findings is that the significant effects in women and men were found in the morning subgroups only. Estradiol, similarly, as testosterone, has a diurnal rhythm in women and men (Bao et al., 2003; Grotzinger et al., 2024). Its diurnal rhythm has been related to diurnal changes in functional connectivity of the brain networks involved in decision-making (Grotzinger et al., 2024). Therefore, this rhythm might amplify the behavioral effects of the hormone at the peak and bottom points in estradiol concentration either during the day or at specific menstrual cycle phases in women.

This aligns with the increasing understanding of a broader chronobiological system which coordinates reproductive, metabolic, cognitive, and motivational functions across time (Joye & Evans, 2022). Some participants in our study were able to choose between their preferred measurement time while the time slots were available, if they fulfilled selection criteria for group similarity. While we did not control for the early vs. late chronotype, it is possible that some participants who were measured in the morning preferred this measurement time due to their chronobiological rhythm, which, in turn, impacted the independent variable’s strength of prediction in our hierarchical models. It is important to highlight that mean estradiol concentrations did not differ between morning and evening women and men groups in our study. Yet future research should investigate whether the behavioral effects might have been influenced by the differences in slope of morning estradiol increase or at what time exactly the participants experienced the peak estradiol concentration, or whether this is an effect of the cross-sectional design of this study.

#### 4.2.4. No testosterone effects

We did not find significant associations between testosterone and moral reasoning parameters. This finding adds to the growing evidence that testosterone may not be reliably associated with moral decision-making, or that any effects are small and only emerge under specific conditions, such as in single-sex samples or as a function of prenatal testosterone exposure, for which the 2D:4D ratio has been used as a proxy (Arnocky et al., 2017; Brannon et al., 2019; Chen et al., 2016; Montoya et al., 2013; Reynolds et al., 2021). This points to a need of hypothesis-driven interaction analysis to assess whether testosterone might be associated with either sex, gender, personality traits or moral inclinations in its effect on moral decision-making.

### 4.3. Personality factors in moral decision-making

In this study, various personality factors were included as predictors of the behavioral parameters in the CNI model. While multiple personality factors emerged as significant predictors in our models, our sensitivity analyses pointed to robust associations between Instrumental Harm and sensitivity to consequences as well as sensitivity to norms in the whole sample, and ARES Behavioral Inhibition and general inaction tendency in women.

#### 4.3.1. Effects of agreement to Instrumental Harm

Agreement to Instrumental Harm was consistently positively associated with the sensitivity to consequences (C parameter) in the whole sample and entire men group. It was also negatively related to sensitivity to norms (N parameter) in the whole sample and in the entire women group.

The Instrumental Harm subscale from the Oxford Utilitarianism Scale (Ambrasė et al., 2025; Kahane et al., 2018) aims to measure whether participants agree to sacrifice a smaller number of individuals to save a larger number of individuals, situations in which moral rightness of the action is measured by better overall consequences (Kahane et al., 2018). In our study, participants who agreed with OUS Instrumental Harm to a higher extent had lower sensitivity to norms. This result mirrors previous findings (Körner et al., 2020). Furthermore, agreement with OUS Instrumental Harm predicted higher sensitivity to consequences. This result contrasts with results from Körner et al. (2020), where the OUS Instrumental Harm subscale was not associated with the sensitivity to consequences in the CNI model, suggesting potential sample effects. In the study by Körner et al. (2020), participants were recruited via MTurk survey platform, were English native speakers and on average older than participants in our study.

The different directions, in which the OUS Instrumental Harm and sensitivity to consequences or norms are related, raise an interesting theoretical issue. Traditionally, utilitarian decision-making, based on consideration of consequences, has been directly contrasted with deontological decision-making, based on norm following, in psychological studies (e.g., Greene et al., 2001). However, it has been argued theoretically that consideration of consequences and norm following are not opposing principles on the same spectrum, but rather two independent moral heuristics (Conway & Gawronski, 2013; Gawronski et al., 2017; Love et al., 2016), which should be clearly separated by the CNI model (Gawronski et al., 2017). Now, the results from our and the previous study (Körner et al., 2020) indicate that moral inclination to agree with Instrumental Harm might be driven by both, higher consideration of consequences or reduced norm following. This suggests that the CNI model might not successfully separate the two moral heuristics, supporting previous CNI critiques regarding the consequences to the model arising from violation of the invariance assumption (Skovgaard-Olsen & Klauer, 2024) and general multinomial processing assumptions (Erdfelder et al., 2009). Future studies should further investigate whether possible interdependence of sensitivity to consequences and sensitivity to norms arises from the architecture of individuals’ moral cognition or from methodological issues within the multinomial processing models.

#### 4.3.2. Effects of Behavioral Inhibition

We found that ARES Behavioral Inhibition negatively predicted inaction in the entire women group and the evening women subgroup. This finding challenges our expectation to find a positive association between ARES Behavioral Inhibition with general inaction tendency. ARES Behavioral Inhibition assesses factors such as anxiousness and frustration intolerance that could reduce individual’s willingness to engage in action. It has been argued that a certain level of disinhibition is necessary to intervene in moral dilemmas and achieve better overall consequences (van den Bos et al., 2011; van den Bos et al., 2009), while ARES Behavioral Inhibition relates to reduced use of instrumental harm (Zhao et al., 2016). However, previous studies did not consider sex/gender differences in these associations, and our results, while unexpected, might suggest that in women ARES Behavioral Inhibition is related to use of other moral reasoning strategies.

## 5. Conclusion

We report that individual biological (sex hormones) and psychological factors (moral inclinations and personality factors) contribute to how individuals arrive at moral decisions differently in women and men. We did not find evidence that women and men differ in their moral reasoning, nor that gender expression would contribute to these differences. Our findings rather reflect the complexity of various influencing factors on moral behavior. Importantly, while women and men did not differ in their moral reasoning in our study, we found distinct predictors contributing to their moral heuristics. It was more difficult for women to make the decisions when imperatives of moral norms did not coincide with the cost-benefit ratio of the proposed moral action. This suggests that while women and men might not differ in how sensitive they are to consequences or norms in moral decision-making, decision difficulty in combination with other biological and personality factors might contribute to previous findings reporting sex/gender differences in frequency of using instrumental harm.

Notably, we found that personality, and specifically, moral inclination towards agreement with Instrumental Harm, is associated with sensitivity to consequences and sensitivity to norms in both women and men. However, the biological influence of gonadal hormones, specifically, estradiol, affected women and men differently – in women, it was positively related to sensitivity to consequences, while in men, negatively related to general inaction tendency. Thus, this study provides first evidence of estradiol’s involvement in moral decision-making. Furthermore, it distinguishes the effects of hormones and personality in one sample, factors that previously have mostly been assessed separately. Finally, it contributes to the still understudied effects of gonadal hormones on value-based decision-making in contexts where outcomes might be negative to the decision-maker or individuals around.

## Notes

### Competing Interest Statement

The authors have declared no competing interest.

### Funding

- Deutsche Forschungsgemeinschaft: DE2319, IRTG 2804

