## Supplemental material for "Sex-specific associations of estradiol and instrumental harm with moral decision-making"

#### Sample characteristics

Supplemental Table 1. Sample characteristics split by measurement time

| Measurement time | Baseline characteristic | n | % |
|---|---|---|---|
| *Morning (7 a.m. – 8 a.m.)* | *Sex/gender* | | |
| | Women | 36 | 53 |
| | Men | 32 | 47 |
| | *Age (years)* | | |
| | 18-20 | 9 | 13 |
| | 21-25 | 41 | 60 |
| | 26-30 | 12 | 17 |
| | 31-35 | 6 | 9 |
| | *Highest education level* | | |
| | General (high school) | 44 | 65 |
| | Bachelor | 19 | 28 |
| | Master | 5 | 7 |
| | Other | 1 | 1 |
| *Evening (3 a.m. – 5 a.m.)* | *Sex/gender* | | |
| | Women | 37 | 64 |
| | Men | 32 | 46 |
| | *Age (years)* | | |
| | 18-20 | 12 | 17 |
| | 21-25 | 44 | 64 |
| | 26-30 | 10 | 15 |
| | 31-35 | 3 | 4 |
| | *Highest education level* | | |
| | General (high school) | 48 | 70 |
| | Bachelor | 16 | 23 |
| | Master | 4 | 6 |
| | Other | 1 | 1 |

### Intercorrelations between predictors

#### Supplemental Table S2. Intercorrelation matrix for the predictors in the study

| | 1. | 2. | 3. | 4. | 5. | 6. | 7. | 8. | 9. | 10. | 11. | 12. | 13. | 14. | 15. |
|---|---|---|---|---|---|---|---|---|---|---|---|---|---|---|---|
| 1. Sex/gender | - | | | | | | | | | | | | | | |
| 2. Measurement time | .007 | - | | | | | | | | | | | | | |
| 3. Femininity | **.43*** ** | -.09 | - | | | | | | | | | | | | |
| 4. Masculinity | **-.41*** ** | .01 | -.11 | - | | | | | | | | | | | |
| 5. Estradiol (log) | **.62*** ** | .02 | **.18* ** | **-.28*** ** | - | | | | | | | | | | |
| 6. Testosterone (log) | **-.97*** ** | -.06 | **-.41*** ** | **.4*** ** | **-.55*** ** | - | | | | | | | | | |
| 7. Progesterone (log) | **.41*** ** | -.14 | .1 | **-.19* ** | **.58*** ** | **-.36*** ** | - | | | | | | | | |
| 8. Cortisol (log) | **-.19* ** | **-.66*** ** | .01 | .08 | **-.24** ** | **.23** ** | -.04 | - | | | | | | | |
| 9. OUS Impartial Beneficence | **.31*** ** | -.04 | **.17* ** | **-.29*** ** | **.25** ** | **-.28*** ** | .07 | .01 | - | | | | | | |
| 10. OUS Instrumental Harm | -.01 | -.004 | -.09 | 0 | -.01 | .01 | -.1 | .03 | .02 | - | | | | | |
| 11. TCI Reward Dependence | **.28*** ** | -.15 | **.49*** ** | **-.19* ** | .16 | **-.25** ** | .03 | -.01 | **.25** ** | -.11 | - | | | | |
| 12. MFQ Care/Harm Foundation | **.22** ** | .06 | **.24** ** | -.07 | **.23** ** | **-.17* ** | **.18* ** | -.06 | **.35*** ** | **-.2* ** | **.28*** ** | - | | | |
| 13. SPF Empthic Concern | **.26** ** | -.09 | **.45*** ** | -.1 | .17 | **-.25** ** | **.2* ** | -.05 | **.26** ** | -.04 | **.4*** ** | **.37*** ** | - | | |
| 14. ARES Behavioural Inhibition | **.24** ** | .06 | .14 | **-.2* ** | .16 | **-.22** ** | .03 | -.12 | .08 | -.03 | **.3*** ** | .05 | **.29*** ** | - | |
| 15. STAI State | **.17* ** | .16 | .09 | **-.29*** ** | .15 | -.17 | .15 | -.13 | .1 | .02 | **.19* ** | -.02 | .1 | **.33*** ** | - |
| 16. BSI Global Symptom Severity | -.11 | .05 | -.02 | -.04 | -.01 | .12 | -.02 | .005 | .06 | .15 | .04 | -.04 | .1 | **.21* ** | **.24** ** |

*** $p \leq 0.001$ (2-tailed), ** $p \leq 0.01$ level (2-tailed), * $p \leq 0.05$ (2-tailed).

#### Sex/gender differences in the CNI model

**Supplemental Table S3. Descriptive statistics for sensitivity to consequences (C parameter) by sex/gender (woman, man) and measurement time (morning, evening)**

| Measure | Women (N = 71) | | Men (N = 64) | |
|---|---|---|---|---|
| | M | SD | M | SD |
| Morning measurement | 0.33 | 0.18 | 0.35 | 0.2 |
| Evening measurement | 0.33 | 0.19 | 0.36 | 0.2 |

**Supplemental Table S4. Two-way ANOVA results for sensitivity to consequences (C parameter)**

| | Sum of squares | df | Mean Square | F | p |
|---|---|---|---|---|---|
| Sex/gender | .035 | 1 | .035 | .92 | .34 |
| Measurement time | .001 | 1 | .001 | .02 | .89 |
| Sex/gender * Measurement time | .002 | 1 | .002 | .04 | .84 |
| Residuals | 4.94 | 131 | .038 | | |

**Supplemental Table S5. Descriptive statistics for sensitivity to norms (N parameter) by sex/gender (woman, man) and measurement time (morning, evening)**

| Measure | Women (N = 71) | | Men (N = 64) | |
|---|---|---|---|---|
| | M | SD | M | SD |
| Morning measurement | 0.77 | 0.26 | 0.66 | 0.29 |
| Evening measurement | 0.56 | 0.32 | 0.59 | 0.31 |

**Supplemental Table S6. Two-way ANOVA results for sensitivity to norms (N parameter)**

| | Sum of squares | df | Mean Square | F | p |
|---|---|---|---|---|---|
| Sex/gender | .06 | 1 | .06 | .67 | .42 |
| Measurement time | .61 | 1 | .61 | 6.5 | .01 |
| Sex/gender * Measurement time | .18 | 1 | .18 | 1.92 | .17 |
| Residuals | 12.38 | 131 | .09 | | |

**Supplemental Table S7. Descriptive statistics for General Inaction/Action tendency (I parameter) by sex/gender (woman, man) and measurement time (morning, evening)**

| Measure | Women (N = 71) | | Men (N = 64) | |
|---|---|---|---|---|
| | M | SD | M | SD |
| Morning measurement | .63 | .4 | .65 | .35 |
| Evening measurement | .59 | .34 | .57 | .35 |

**Supplemental Table S8. Two-way ANOVA results for General Inaction/Action tendency (I parameter)**

| | Sum of squares | df | Mean Square | F | p |
|---|---|---|---|---|---|
| **Sex/gender** | .00006 | 1 | .00006 | 0 | .98 |
| **Measurement time** | .12 | 1 | .12 | .95 | .33 |
| **Sex/gender * Measurement time** | .01 | 1 | .01 | .1 | .75 |
| **Residuals** | 16.92 | 131 | .13 | | |

#### Diagnostics of linear regressions predicting sensitivity to consequences (C parameter)

Diagnostics of linear regressions predicting the C parameter (Table 1) indicated deviations from normality in the entire men group (Step 4) and the morning men group (Step 1) models and violations of homoscedasticity in the evening men group (Step 2) and the entire women group (Step 4) models, as assessed by formal statistical tests. However, visual inspection of Q-Q plots and residual plots suggested that these deviations were minor and did not indicate substantial violations of model assumptions. Influential data points have been observed in the morning men's (Step 4) and women's group (Step 4). However, removal of these observations did not eliminate significant effects of regression models. In morning men's group, after elimination of influential data points masculinity became a significant predictor (B = 0.1, t = 2.33, p = 0.03).

**Supplemental Table S9.** Diagnostics of linear regressions predicting the C parameter. The normality of residuals was investigated using the Shapiro-Wilk test, and the homoscedasticity was evaluated using the Breusch-Pagan test. The highest Cook's distance value is reported for each regression model.

| | **C parameter** | | | |
|---|---|---|---|---|
| | **Step 1** | **Step 2** | **Step 3** | **Step 4** |
| *Whole sample* | | | | |
| Normality | 0.98 (p = 0.12) | 0.99 (p = 0.15) | 0.98 (p = 0.11) | 0.99 (p = 0.17) |
| Homoscedasticity | 0.54 (p = 0.97) | 7.28 (p = 0.51) | 6.04 (p = 0.53) | 17.65 (p = 0.09) |
| Cook's distance | 0.08 | 0.069 | 0.086 | 0.093 |
| *Entire men group* | | | | |
| Normality | 0.97 (p = 0.11) | 0.97 (p = 0.1) | 0.97 (p = 0.11) | **0.96 (p = 0.04)** |
| Homoscedasticity | 1.49 (p = 0.47) | 5.94 (p = 0.43) | 1.45 (p = 0.92) | 7.08 (p = 0.63) |
| Cook's distance | 0.079 | 0.082 | 0.183 | 0.207 |
| *Morning men group* | | | | |
| Normality | **0.93 (p = 0.04)** | 0.95 (p = 0.15) | 0.97 (p = 0.62) | 0.97 (p = 0.59) |
| Homoscedasticity | < 0.001 (p = 1) | 7.04 (p = 0.22) | 7.92 (p = 0.09) | 14.53 (p = 0.07) |
| Cook's distance | 0.17 | 0.27 | 0.387 | **0.533** |
| *Evening men group* | | | | |
| Normality | 0.97 (p = 0.57) | 0.96 (p = 0.36) | 0.98 (p = 0.81) | 0.96 (p = 0.23) |
| Homoscedasticity | 2.8 (p = 0.09) | **14.5 (p = 0.01)** | 3.42 (p = 0.49) | 8.06 (p = 0.43) |
| Cook's distance | 0.156 | 0.284 | 0.25 | 0.186 |
| *Entire women group* | | | | |
| Normality | 0.98 (p = 0.4) | 0.98 (p = 0.34) | 0.99 (p = 0.57) | 0.99 (p = 0.66) |
| Homoscedasticity | 0.51 (p = 0.77) | 6.97 (p = 0.32) | 7.34 (p = 0.2) | **21.36 (p = 0.01)** |
| Cook's distance | 0.236 | 0.151 | 0.157 | 0.128 |
| *Morning women group* | | | | |
| Normality | 0.98 (p = 0.9) | 0.97 (p = 0.52) | 0.98 (p = 0.79) | 0.97 (p = 0.62) |
| Homoscedasticity | 0.67 (p = 0.41) | 2.25 (p = 0.81) | 1.53 (p = 0.82) | 6.45 (p = 0.6) |
| Cook's distance | 0.136 | 0.361 | 0.182 | **0.567** |
| *Evening women group* | | | | |
| Normality | 0.97 (p = 0.37) | 0.98 (p = 0.6) | 0.98 (p = 0.88) | 0.98 (p = 0.62) |
| Homoscedasticity | 0.31 (p = 0.58) | 2.49 (p = 0.78) | 2.02 (p = 0.73) | 8.71 (p = 0.37) |
| Cook's distance | 0.371 | 0.203 | 0.245 | 0.325 |

#### Diagnostics of linear regressions predicting sensitivity to norms (N parameter)

Diagnostics of linear regressions predicting the N parameter (Table 2) indicated multiple deviations from normality. However, inspection of the Q-Q plots did not indicate strong evidence of non-normality in any of the models. Influential data points have been observed in the evening men's group (Steps 2 and 4). However, removal of these observations did not meaningfully affect the regression models.

**Supplemental Table S10.** Diagnostics of linear regressions predicting the N parameter. The normality of residuals was investigated using the Shapiro-Wilk test, and the homoscedasticity was evaluated using the Breusch-Pagan test. The highest Cook's distance value is reported for each regression model.

| | N parameter | | | |
|---|---|---|---|---|
| | **Step 1** | **Step 2** | **Step 3** | **Step 4** |
| *Whole sample* | | | | |
| Normality | **0.95 (p < 0.001)** | **0.95 (p < 0.001)** | **0.96 (p < 0.001)** | **0.96 (p < 0.001)** |
| Homoscedasticity | 5.83 (p = 0.21) | 10.15 (p = 0.25) | 9.85 (p = 0.28) | 13.1 (p = 0.36) |
| Cook's distance | 0.083 | 0.102 | 0.067 | 0.075 |
| *Entire men group* | | | | |
| Normality | **0.92 (p < 0.001)** | **0.93 (p = 0.001)** | **0.94 (p = 0.004)** | **0.94 (p = 0.003)** |
| Homoscedasticity | 1.84 (p = 0.4) | 4.44 (p = 0.62) | 5.76 (p = 0.45) | 8.08 (p = 0.62) |
| Cook's distance | 0.126 | 0.312 | 0.127 | 0.156 |
| *Morning men group* | | | | |
| Normality | **0.9 (p = 0.007)** | **0.9 (p = 0.008)** | **0.89 (p = 0.004)** | **0.92 (p = 0.017)** |
| Homoscedasticity | 0.04 (p = 0.84) | 2.58 (p = 0.76) | 5.4 (p = 0.37) | 10.85 (p = 0.29) |
| Cook's distance | 0.115 | 0.114 | 0.166 | 0.255 |
| *Evening men group* | | | | |
| Normality | **0.91 (p = 0.014)** | 0.96 (p = 0.26) | **0.92 (p = 0.02)** | **0.93 (p = 0.03)** |
| Homoscedasticity | 1.36 (p = 0.24) | 7.24 (p = 0.2) | 1.52 (p = 0.91) | 4.16 (p = 0.9) |
| Cook's distance | 0.424 | **1.45** | 0.27 | **1.18** |
| *Entire women group* | | | | |
| Normality | **0.94 (p = 0.003)** | **0.95 (p = 0.008)** | **0.95 (p = 0.007)** | **0.94 (p = 0.002)** |
| Homoscedasticity | 2 (p = 0.37) | 4.88 (p = 0.56) | 7.41 (p = 0.28) | 11.63 (p = 0.31) |
| Cook's distance | 0.133 | 0.125 | 0.14 | 0.123 |
| *Morning women group* | | | | |
| Normality | **0.87 (p < 0.001)** | **0.94 (p = 0.06)** | **0.9 (p = 0.006)** | **0.93 (p = 0.04)** |
| Homoscedasticity | 1.3 (p = 0.25) | 4.1 (p = 0.53) | 7.25 (p = 0.2) | 6.37 (p = 0.7) |
| Cook's distance | 0.232 | 0.135 | 0.346 | 0.228 |
| *Evening women group* | | | | |
| Normality | **0.93 (p = 0.02)** | 0.96 (p = 0.27) | 0.97 (p = 0.32) | 0.95 (p = 0.08) |
| Homoscedasticity | 0.15 (p = 0.7) | 6.93 (p = 0.23) | 4.5 (p = 0.48) | 9.11 (p = 0.43) |
| Cook's distance | 0.214 | 0.231 | 0.203 | 0.312 |

#### Diagnostics of linear regressions predicting general inaction/action tendency (I parameter)

Diagnostics of linear regressions predicting the I parameter (Table 3) indicated deviations from normality in several models. Visual inspection of Q–Q plots in the whole sample indicated substantial deviations from normality, with pronounced skewness in residuals (especially in Steps 1 and 2). Similar deviations based on Q-Q plots were observed in Steps 1 and 2 of the entire men group, the morning men group, and Step 1 of the evening men group. Substantial deviations according to the inspection of Q-Q plots were observed in Step 1 of the entire women's group, Steps 1 and 2 in the morning women's group, and Step 1 in the evening women's group.

Inspection of residuals versus fitted values plots indicated limited evidence of heteroscedasticity, with notable deviations observed only in Step 2 of the full men sample. In other models, visual inspection did not suggest meaningful violations of homoscedasticity.

Influential data points have been observed in the morning men's group (Steps 2, 3 and 4). After removing the influential observation in Step 2, estradiol remained a significant predictor (B = 1.19, SE = 0.22, t = 5.44, $p < 0.001$), and masculinity became a significant negative predictor of the I parameter (B = -0.18, SE = 0.07, t = -2.5, p = 0.02). Removal of the influential observation in Step 3 did not meaningfully affect the model. After removing two influential observations in Step 4, estradiol remained a significant predictor (B = 1.27, SE = 0.2, t = 6.24, $p < 0.001$), masculinity became a significant negative predictor (B = 0.15, SE = 0.07, t = -2.14, p = 0.04), STAI state became a significant positive predictors (B = 0.01, SE = 0.005, t = 2.37, p = 0.03), and anxiousness became a significant negative predictor (B = -0.45, SE = 0.19, t = -2.43, p = 0.02).

Influential data points have also been observed in the morning women's group (Steps 2 and 4). However, removing these observations did not meaningfully change the regression models.

**Table 3.** Diagnostics of linear regressions predicting the I parameter. The normality of residuals was investigated using the Shapiro-Wilk test, and the homoscedasticity was evaluated using the Breusch-Pagan test. The highest Cook's distance value is reported for each regression model.

| | **I parameter** | | | |
|---|---|---|---|---|
| | **Step 1** | **Step 2** | **Step 3** | **Step 4** |
| *Whole sample* | | | | |
| Normality | **0.9 (p < 0.001)** | **0.9 (p < 0.001)** | **0.94 (p < 0.001)** | **0.94 (p < 0.001)** |
| Homoscedasticity | 1.59 (p = 0.81) | 2.54 (p = 0.96) | 6.26 (p = 0.51) | 6.34 (p = 0.85) |
| Cook's distance | 0.053 | 0.134 | 0.122 | 0.095 |
| *Entire men group* | | | | |
| Normality | **0.91 (p < 0.001)** | 0.97 (0.12) | **0.91 (p < 0.001)** | 0.98 (p = 0.25) |
| Homoscedasticity | 0.27 (p = 0.87) | **16.3 (p = 0.012)** | 1.34 (p = 0.93) | 15.26 (p = 0.08) |
| Cook's distance | 0.113 | 0.323 | 0.172 | 0.25 |
| *Morning men group* | | | | |
| Normality | **0.86 (p < 0.001)** | 0.98 (p = 0.72) | **0.88 (p = 0.002)** | 0.99 (p = 0.95) |
| Homoscedasticity | 0.1 (p = 0.76) | **13.7 (p = 0.02)** | 1.5 (p = 0.83) | **16.6 (p = 0.03)** |
| Cook's distance | 0.291 | **0.686** | **0.524** | **0.609** |
| *Evening men group* | | | | |
| Normality | **0.9 (p = 0.005)** | 0.94 (p = 0.09) | 0.94 (p = 0.08) | 0.96 (p = 0.23) |
| Homoscedasticity | 1.49 (p = 0.22) | 4.51 (p = 0.48) | 3.14 (p = 0.53) | 6.44 (p = 0.6) |
| Cook's distance | 0.29 | 0.421 | 0.23 | 0.463 |
| *Entire women group* | | | | |

| | | | | |
|---|---|---|---|---|
| Normality | **0.89 (p < 0.001)** | **0.93 (p < 0.001)** | **0.96 (p = 0.03)** | 0.97 (p = 0.11) |
| Homoscedasticity | 1.97 (p = 0.37) | 9.35 (p = 0.15) | 10.06 (p = 0.07) | 8.99 (p = 0.44) |
| Cook's distance | 0.111 | 0.131 | 0.108 | 0.085 |
| *Morning women group* | | | | |
| Normality | **0.83 (p < 0.001)** | **0.84 (p < 0.001)** | 0.94 (p = 0.08) | 0.94 (p = 0.06) |
| Homoscedasticity | 0.53 (p = 0.47) | 5.7 (p = 0.34) | 1.43 (p = 0.84) | 9.87 (p = 0.27) |
| Cook's distance | 0.393 | **1.69** | 0.216 | **1.74** |
| *Evening women group* | | | | |
| Normality | **0.92 (p = 0.01)** | 0.97 (p = 0.49) | 0.97 (p = 0.49) | 0.96 (p = 0.26) |
| Homoscedasticity | 0.01 (p = 0.91) | 13.55 (p = 0.02) | 1.52 (p = 0.82) | 3.25 (p = 0.92) |
| Cook's distance | 0.16 | 0.477 | 0.17 | 0.379 |

#### Sensitivity power analysis

To evaluate the robustness of the regression findings, a sensitivity power analysis using G*Power 3.1.9.7 (Faul et al., 2009) was conducted for those models that revealed significant predictors. The required effect sizes for each sample and each Step were calculated assuming α = 0.05, power = 0.8, tested predictor = 1, and appropriate sample sizes and total numbers of predictors. The minimal detectable effect size ($f^2$) for each sample and its Step, along with predictors' partial $f^2$, are provided in Table 4.

**Supplemental Table S12.**

| Parameter and sample | Step | Minimal $f^2$ | Predictors' partial $f^2$ |
|---|---|---|---|
| **Sensitivity to consequences** | | | |
| *Whole sample* | | | |
| | 3 | 0.059 | Instrumental harm ($f^2$ = 0.148)*, psychopathy ($f^2$ = 0.045) |
| | 4 | 0.06 | Instrumental harm ($f^2$ = 0.151)*, psychopathy ($f^2$ = 0.046) |
| *Entire men group* | | | |
| | 3 | 0.127 | Instrumental harm ($f^2$ = 0.173)* |
| | 4 | 0.129 | Instrumental harm ($f^2$ = 0.173)* |
| *Morning men group* | | | |
| | 3 | 0.265 | Instrumental harm ($f^2$ = 0.384)*, Impartial Beneficence ($f^2$ = 0.18) |
| | 4 | 0.269 | Instrumental harm ($f^2$ = 0.436)* |
| *Evening men group* | | | |
| | 4 | 0.278 | Reward dependency ($f^2$ = 0.256) |
| *Entire women group* | | | |
| | 3 | 0.114 | Instrumental harm ($f^2$ = 0.103) |
| | 4 | 0.116 | Instrumental harm ($f^2$ = 0.106) |
| *Morning women group* | | | |
| | 2 | 0.26 | Estradiol ($f^2$ = 0.223), progesterone ($f^2$ = 0.246) |
| | 4 | 0.26 | Estradiol ($f^2$ = 0.374)*, progesterone ($f^2$ = 0.135) |
| **Sensitivity to norms** | | | |
| *Whole sample* | | | |
| | 1 | 0.059 | Time ($f^2$ = 0.05) |
| | 3 | 0.059 | Time ($f^2$ = 0.057), Instrumental harm ($f^2$ = 0.075)* |
| | 4 | 0.06 | Instrumental harm ($f^2$ = 0.025) |
| *Morning men group* | | | |
| | 3 | 0.265 | Instrumental harm ($f^2$ = 0.189), Care/Harm foundation ($f^2$ = 0.204) |
| | 4 | 0.269 | Instrumental harm ($f^2$ = 0.261), Care/Harm foundation ($f^2$ = 0.223) |
| *Entire women sample* | | | |
| | 1 | 0.114 | Time ($f^2$ = 0.126)* |
| | 3 | 0.116 | Time ($f^2$ = 0.182)* |
| | 4 | 0.118 | Instrumental harm ($f^2$ = 0.085) |
| **General Inaction/Action tendency** | | | |
| *Entire men group* | | | |

| | | | |
|---|---|---|---|
| | 2 | 0.129 | Estradiol ($f^2$ = 0.104) |
| | 4 | 0.129 | Estradiol ($f^2$ = 0.111) |
| *Morning men group* | | | |
| | 2 | 0.265 | Estradiol ($f^2$ = 0.634)* |
| | 4 | 0.267 | Estradiol ($f^2$ = 0.631)* |
| *Entire women group* | | | |
| | 3 | 0.114 | Behavioural Inhibition ($f^2$ = 0.118)*, State Anxiety ($f^2$ = 0.078) |
| | 4 | 0.116 | Behavioural Inhibition ($f^2$ = 0.135)*, State Anxiety ($f^2$ = 0.092) |
| *Morning women group* | | | |
| | 3 | 0.247 | Behavioural Inhibition ($f^2$ = 0.145), State Anxiety ($f^2$ = 0.166) |
| *Evening women group* | | | |
| | 3 | 0.226 | Behavioural Inhibition ($f^2$ = 0.308)*, Anxiousness ($f^2$ = 0.298)* |
| | 4 | 0.228 | Behavioural Inhibition ($f^2$ = 0.395)*, Anxiousness ($f^2$ = 0.267)* |

* Predictor's effect size exceeded the detection threshold

#### Sex/gender differences in decision difficulty

**Supplemental Table S13. Descriptive statistics for decision difficulty prospective scenarios when benefits are greater than the costs by biological sex (woman, man) and measurement time (morning, evening)**

| Measure | Women | | Men | |
|---|---|---|---|---|
| | M | SD | M | SD |
| Morning measurement | 4.27 | 0.97 | 3.94 | 1 |
| Evening measurement | 4.45 | 0.96 | 4.02 | 1.2 |

**Supplemental Table S14. Two-way ANOVA results for decision difficulty prospective scenarios when benefits are greater than the costs by biological sex (woman, man) and measurement time (morning, evening)**

| | Sum of squares | df | Mean Square | F | p |
|---|---|---|---|---|---|
| Biological sex | 4.97 | 1 | 4.97 | 4.76 | .03* |
| Measurement time | .58 | 1 | .58 | .55 | .46 |
| Biological sex * Measurement time | .07 | 1 | .07 | .07 | .79 |
| Residuals | 137.83 | 132 | 1.04 | | |

**Supplemental Table S15. Descriptive statistics for decision difficulty prospective scenarios when benefits are smaller than the costs by biological sex (woman, man) and measurement time (morning, evening)**

| Measure | Women | | Men | |
|---|---|---|---|---|
| | M | SD | M | SD |
| Morning measurement | 2.08 | .7 | 2.19 | .72 |
| Evening measurement | 2.28 | .73 | 2.19 | .74 |

**Supplemental Table S16. Two-way ANOVA results for decision difficulty prospective scenarios when benefits are smaller than the costs by biological sex (woman, man) and measurement time (morning, evening)**

| | Sum of squares | df | Mean Square | F | p |
|---|---|---|---|---|---|
| Biological sex | .006 | 1 | .006 | .01 | .92 |
| Measurement time | .31 | 1 | .31 | .58 | .45 |
| Biological sex * Measurement time | .33 | 1 | .33 | .62 | .43 |
| Residuals | 69.34 | 132 | .53 | | |

**Supplemental Table S17. Descriptive statistics for decision difficulty prespective scenarios when benefits are greater than the costs by biological sex (woman, man) and measurement time (morning, evening)**

| Measure | Women | | Men | |
|---|---|---|---|---|
| | M | SD | M | SD |
| Morning measurement | 2.68 | .9 | 2.73 | .61 |
| Evening measurement | 2.91 | .83 | 2.57 | .9 |

**Supplemental Table S18. Two-way ANOVA results for decision difficulty prespective scenarios when benefits are greater than the costs by biological sex (woman, man) and measurement time (morning, evening)**

| | Sum of squares | df | Mean Square | F | p |
|---|---|---|---|---|---|
| Biological sex | .79 | 1 | .79 | 1.18 | .28 |
| Measurement time | .95 | 1 | .95 | .07 | .79 |
| Biological sex * Measurement time | 1.33 | 1 | 1.33 | 1.97 | .16 |
| Residuals | 88.84 | 132 | .67 | | |

**Supplemental Table S19. Descriptive statistics for decision difficulty prespective scenarios when benefits are smaller than the costs by biological sex (woman, man) and measurement time (morning, evening)**

| Measure | Women | | Men | |
|---|---|---|---|---|
| | M | SD | M | SD |
| Morning measurement | 4.46 | .94 | 4.11 | 1.15 |
| Evening measurement | 4.58 | 1.05 | 4.06 | 1.31 |

**Supplemental Table S20. Two-way ANOVA results for decision difficulty prespective scenarios when benefits are smaller than the costs by biological sex (woman, man) and measurement time (morning, evening)**

| | Sum of squares | df | Mean Square | F | p |
|---|---|---|---|---|---|
| Biological sex | 6.41 | 1 | 6.41 | 5.18 | .02* |
| Measurement time | .04 | 1 | .04 | .03 | .86 |
| Biological sex * Measurement time | .24 | 1 | .24 | .19 | .66 |
| Residuals | 163.33 | 132 | 1.24 | | |
